# Morphogenesis-on-chip: A minimal in vitro assay for cell intercalation highlights the importance of interfacial tension and migratory forces

**DOI:** 10.1101/2025.06.30.662274

**Authors:** Artur Ruppel, Vladimir Misiak, Sadjad Arzash, Daniel Selma Herrador, Thomas Boudou, Lisa Manning, François Fagotto, Martial Balland

## Abstract

Cell intercalation, the dynamic exchange of cellular neighbors, is fundamental to embryonic morphogenesis, tissue homeostasis, and wound healing. Despite extensive study in complex tissues, the minimal mechanical requirements driving intercalation remain poorly understood due to confounding tissue level interactions. Here, we present a novel morphogenesis-on-chip assay utilizing micropatterned cell quadruplets. This system isolates the elementary unit of intercalation while enabling quantitative force and shape measurements. Cross-shaped micropatterns generate stable four cell configurations in MDCK epithelial cells. Surprisingly, these cells spontaneously undergo T1 transitions autonomously. We combined live imaging with force inference and traction force microscopy, which revealed that intercalation emerges from two distinct mechanisms: interfacial tension dynamics and differential cell migration. Specifically, we show a correlation between central junction shrinkage and increased relative tension. Similarly, we show a correlation between central junction shrinkage and migratory forces. We successfully adapted the assay to Xenopus mesoderm cells, revealing conserved mechanical principles across cell types. Furthermore, experimentally derived effective energy landscapes closely match theoretical vertex model predictions, and suggest a dominant role for migratory forces in driving intercalation. This confirms that our minimal system recapitulates the fundamental physics of intercalation. This approach provides the first quantitative framework for studying intercalation mechanics in isolation and establishes a versatile platform for investigating morphogenetic processes.

**Graphical Abstract:** 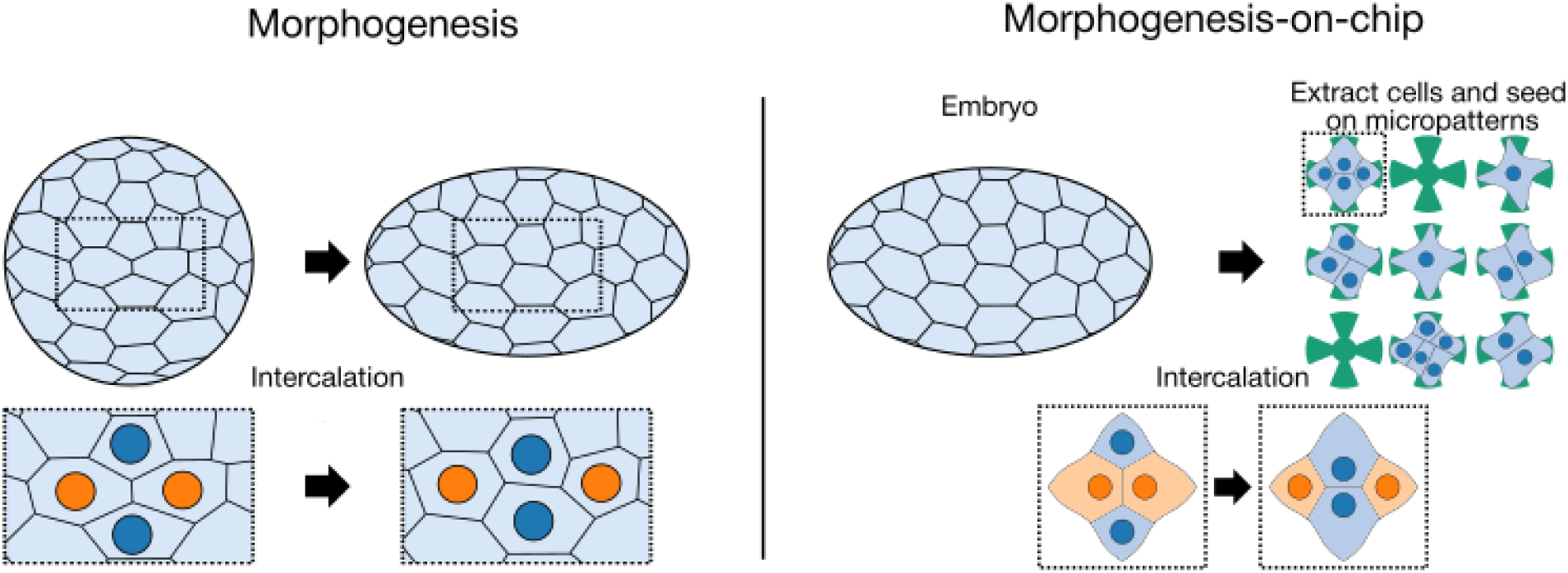

## 1 Introduction

Animal tissues have the remarkable capacity to undergo extensive remodeling while remaining structurally coherent. At the core of this capacity is the ability of cells to dynamically exchange neighbours, a process called cell intercalation [1, 2]. This fundamental cellular behavior is essential for embryonic morphogenesis, tissue homeostasis, regeneration, wound healing, and can be co-opted during cancer progression [2, 3]. While intercalation in living tissues is fundamentally driven by active, cell-generated forces, these rearrangements can also be influenced and induced by external forces, distinguishing them from passive materials like foams that rearrange solely in response to external stimuli. [4]. By regulating intercalation, cells effectively control tissue rheology, with higher intercalation rates typically increasing fluidity [5, 2]. Furthermore, when intercalations are directionally biased, they can generate dramatic large-scale tissue transformations. Radial intercalation drives tissue thinning and spreading, while medial-lateral intercalation enables tissue elongation [6, 7, 8]. In two-dimensional tissues, these complex rearrangements can be decomposed into fundamental topological events called T1 transitions, where two initially contacting cells disengage while two adjacent cells establish a new contact, effectively exchanging neighbors [9].

Two major cellular processes are at the basis of cell intercalation: Contractility of the actomyosin cortex along cell-cell interfaces, and cell motility, which relies on protrusive activity [10, 11, 12, 13, 14]. Cortical contractility can lead to shrinkage of an existing contact, favouring the encounter of other adjacent cells. Protrusions allow cells to crawl on extracellular matrix substrate and on other cells, thus contributing to reposition cells relative to each other. Here intercalation can be driven either by cells moving toward each other, favoring formation of a new contact, or by cells moving away, stimulating disruption of a contact, thus indirectly allowing two other cells to get into contact [15]. These mechanisms can function independently or cooperatively depending on the tissue context and cell type. Increased tension at the shrinking contact appears to dominate T1 transitions in epithelial monolayers such as in Drosophila [11, 16]. Intercalation based on migration was first described for vertebrate mesoderm, a prototype of coherent mesenchymal tissue [17, 18, 14]. It was later shown that polarized myosin-dependent contact shrinkage was also operating during convergent-extension in the mesoderm [19]. Reciprocally, the basal side of epithelial cells also displays protrusive activity and motility, which also contributes to intercalation in epithelial monolayers [20].

Despite the wealth of information gathered over the past two decades, the phenomenon of intercalation and its various flavors remain incompletely understood, both in terms of cellular and molecular mechanisms and physical properties. Open questions include the actual relative contributions of the contractile and migratory mechanisms and their potential crosstalk, the nature, impact and regulation of spontaneous tension fluctuations, and, in the case of oriented intercalations, the interplay between stochastic and cue-oriented reactions and/or persistent motility.

Most current knowledge has come from studies on whole tissues, often within the context of whole embryos. Teasing apart the many variables in such complex systems remains a daunting challenge. Forces are primarily indirectly inferred from imaging data or from laser ablation [21, 22]. Furthermore, these multicellular ensembles behave as integrated systems, with physical forces and molecular information spreading over distance, and individual intercalation events influencing each other [23, 24]. Together, these limitations make it difficult to isolate the minimal mechanical requirements for intercalation.

To complement these existing approaches, there is a pressing need for simplified experimental systems that can replicate essential features of intercalation while allowing for precise measurements and controlled perturbations. Here, we introduce a “morphogenesis-on-chip” platform that reduces tissue rearrangement to its fundamental unit: a four-cell quadruplet. We demonstrate that these minimalistic tissues, formed from either epithelial MDCK or primary Xenopus mesoderm cells, remarkably undergo spontaneous and autonomous cell intercalation. By combining quantitative live imaging with traction force microscopy and force inference, we dissect the contributions of the two key mechanical drivers: interfacial tension dynamics and cell migratory forces. Our results reveal a conserved mechanism where junction shrinkage directly correlates with both increased relative tension at cell-cell contacts and coordinated migratory activity on the substrate. Furthermore, we derive the first experimentally-measured energy landscapes for intercalation, which closely match theoretical vertex model predictions and suggest a dominant role for migratory forces in overcoming the transition barrier. Taken together, this work establishes a powerful and versatile framework for dissecting the fundamental physics of morphogenesis in a precisely controlled environment, uncoupled from the complexity of a whole tissue.

## 2 Results

### 2.1 Cross-shaped micropatterns induce reproducible MDCK cell quadruplets that undergo spontaneous intercalation

The design of the micropattern was guided by previous work demonstrating that cell-cell junctions tend to stabilize over regions devoid of extracellular matrix (ECM) proteins, as demonstrated in cell doublets [25, 26]. We systematically explored various geometries to achieve four-cell quadruplets with reproducible morphology. Initial designs, such as simple square patterns, resulted in cells rotating around each other (Movie S1). Other configurations, like a corner and circle pattern, did not provide sufficiently connected spreading area for cells to reliably encounter each other (Movie S2). A semi-cross pattern led to interactions only between cells aligned along the ECM lines (Movie S3). Ultimately, the full cross-shaped micropattern (Movie S4) proved most effective, consistently promoting the formation of reproducible quadruplet geometries that remained stable for hours in MDCK cells, confirming the principle of junction stabilization over non-ECM areas for these four-cell arrangements.

We thus chose this cross-shaped micropattern to produce a minimal cell configuration that would allow to study intercalation in a controlled environment. On this pattern, MDCK cells formed stable quadruplets, mimicking the basic tiling observed in tissues (Figure 1A). This confined configuration ensured a stereotypical arrangement where all four cells were laterally in contact with each other, with two opposite cells forming a central cell-cell adhesive interface, which we will hereafter refer to as central junction. Imaging of filamentous actin (LifeAct) revealed robust alignment of cell boundaries along the edges of the pattern and consistent positioning of the central junction. The reproducibility of this configuration was evident from averaging LifeAct signal of multiple quadruplets (Figure 1B).

**Figure 1:**
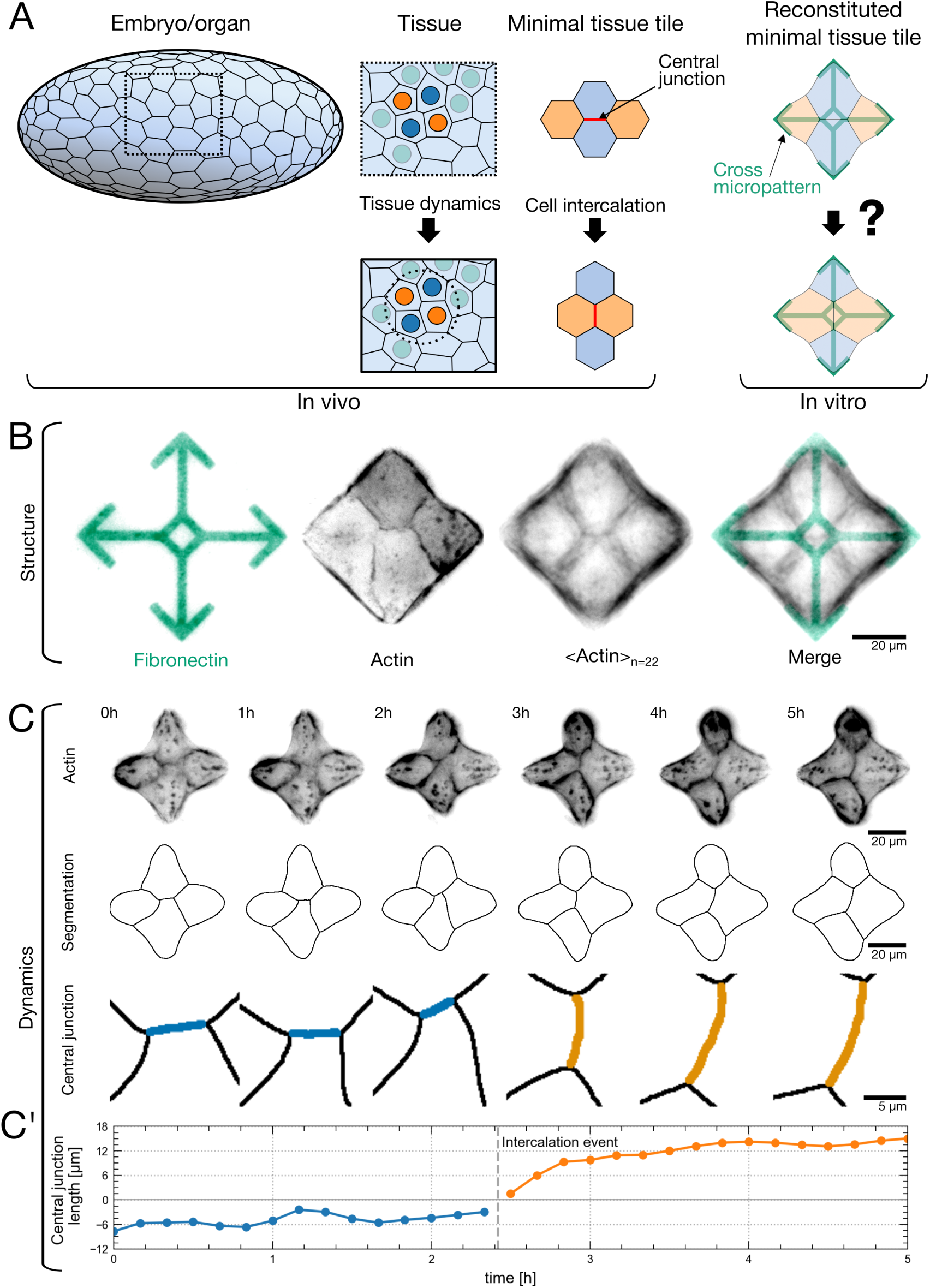
Cross-shaped micropatterns induce reproducible MDCK cell quadruplets that undergo spontaneous intercalation. (A) Schematic representation of a tissue undergoing dynamic cell rearrangement during morphogenesis, driven by cell-cell intercalations. In the simple case of a cell monolayer, a minimal tissue unit can be described as a cell quadruplet, with two cells contributing to a central contact. Intercalation involves the loss of the central contact and establishment of a new contact by the two other cells. This basic change in topography is called a T1 transition. The two schematics on the right illustrate our in vitro reconstitution of this elementary tissue tile. (B) The cross-shaped fibronectin-coated micropattern (green) ensures consistent quadruplet configurations, as shown by live detection of filamentous actin (far-red LifeAct shown in grayscale) of a representative quadruplet. Averaged LifeAct intensity image of 22 quadruplets showing reproducible cell positioning and actin organization, with the final panel showing this average superimposed with the micropattern. (C) Example of spontaneous intercalation in a MDCK quadruplet. Selected frames from a time lapse movie showing: LifeAct images (top row); skeletonized segmentations (second row); enlarged view highlighting the T1 transition, with the original central junction in blue and the new central junction in orange (third row). (C’) Central junction length over time. Negative lengths denote the initial junction state before any intercalation event, and any following intercalation event changes the sign. This process is illustrated in Movie S5 and Movie S6.

**Figure 1, Supplementary Data.**
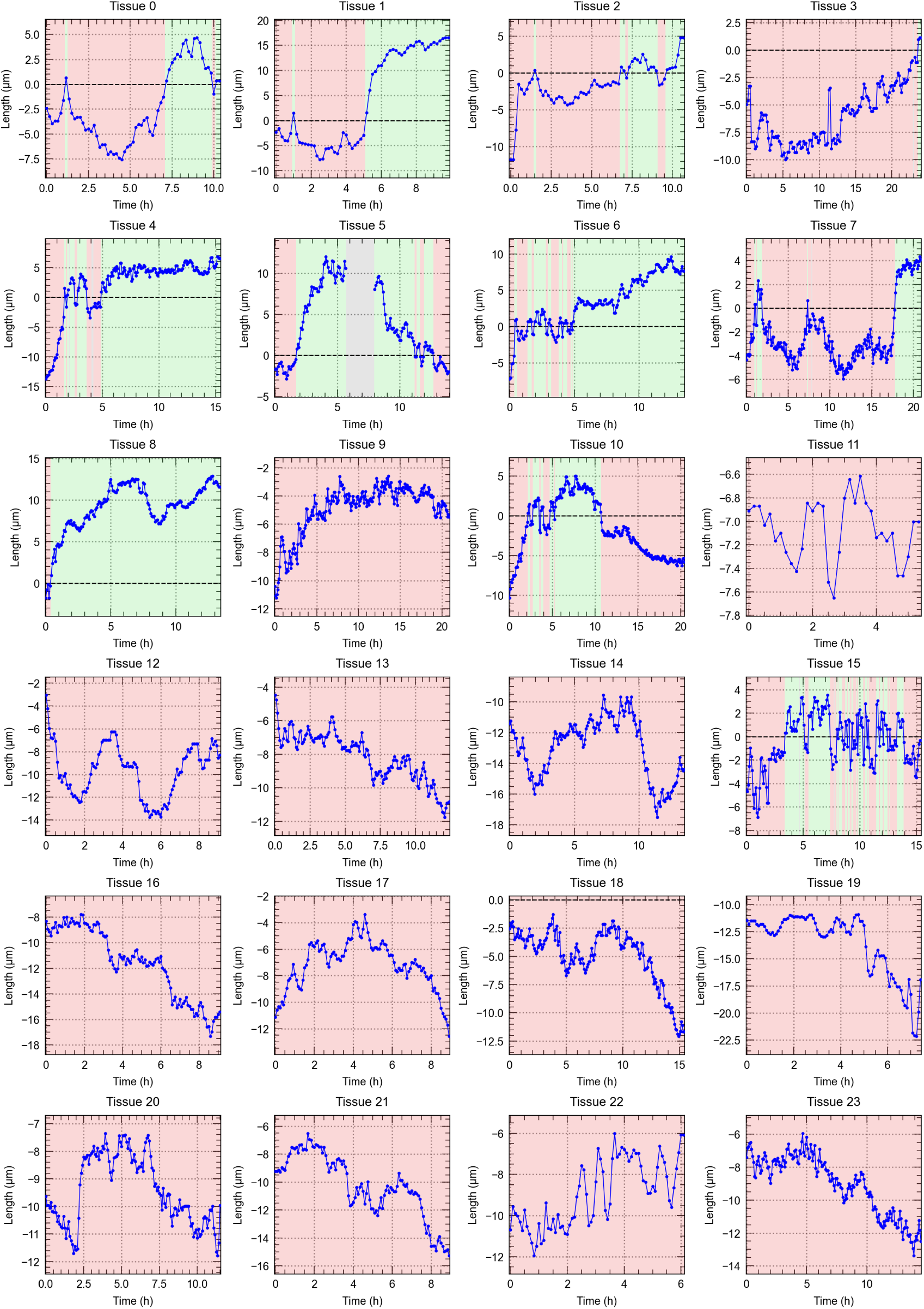
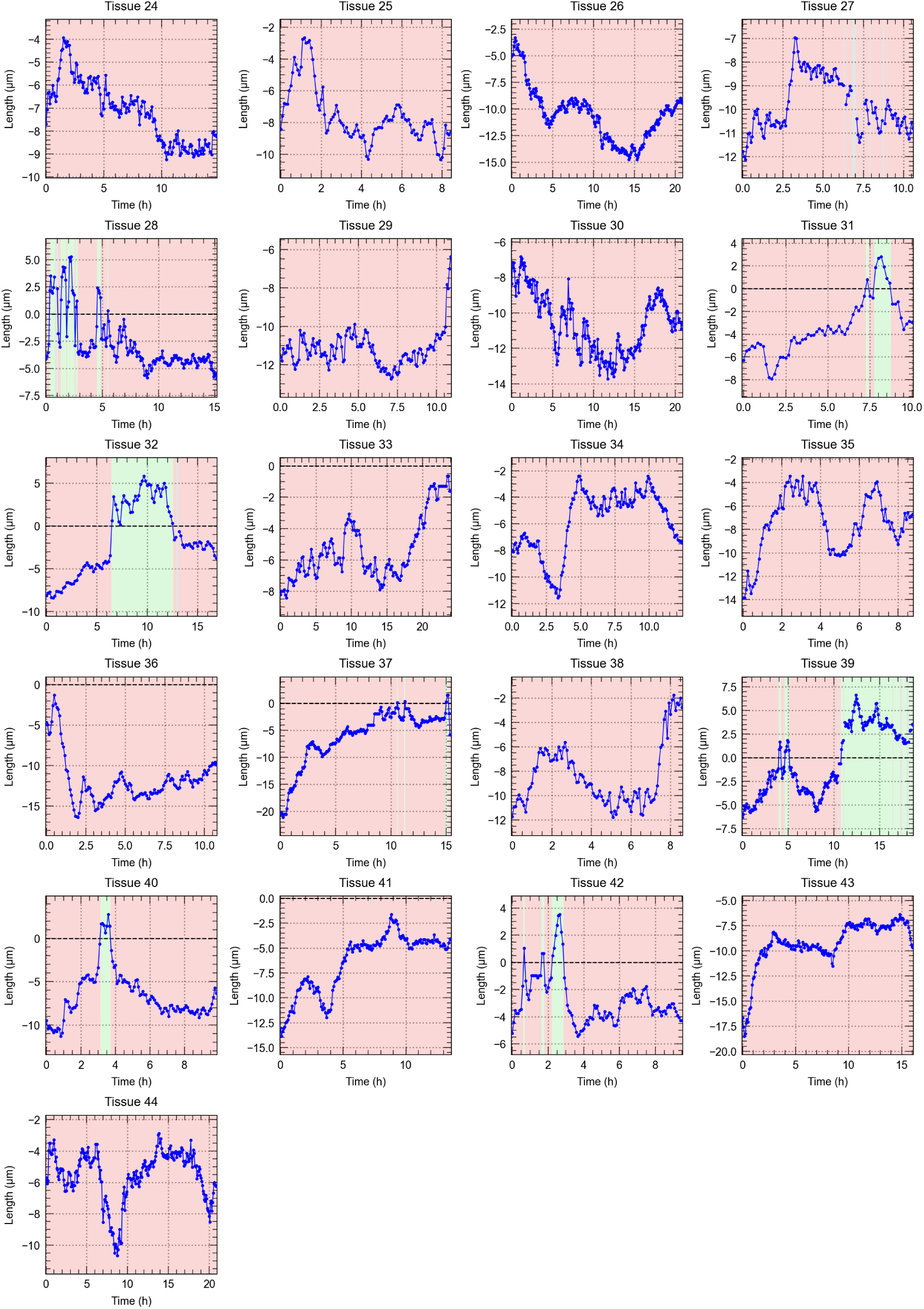
Central junction dynamics and intercalation events in MDCK quadruplets. Signed central junction length over time for individual quadruplets. Negative lengths denote the initial junction state before any intercalation event, and any following intercalation event changes the sign.

Despite the constraints imposed by the micropattern, MDCK cells underwent spontaneous intercalations, with the typical topography of T1 transitions, characterized by the shrinkage of the central junction, its disappearance and the concomitant appearance of a new junction perpendicular to the initial one (Figure 1C, Movie S5, Movie S6). To analyze these experiments we developed napariCellFlow, a napari plugin, for cell segmentation, cell tracking, edge and intercalation detection, and junction length measurement (see Materials and Methods for details). The graph of Figure 1C’ represents the variations in length of the central junction over time, showing typical fluctuations until a phase of shrinkage led to its abrupt disruption and formation of the new junction, which subsequently expanded (Figure 1C-C’, Movie S5, Movie S6). These observations were consistent across multiple replicates, confirming the system’s ability to reproduce the dynamic rearrangements observed in vivo. 17 out of 45 (38%) quadruplets showed at least one intercalation over the imaging window of on average about 12 h (Figure 1, Supplementary Data), many of them intercalating multiple times. Thus, a minimal cell quadruplet could autonomously undergo intercalation with a surprisingly high frequency considering the high stability of the configuration and the absence of instructive cues.

The behavior of the central junction of the quadruplets was highly variable: some junctions maintained relatively stable lengths for hours, others exhibited high variability without intercalation, some intercalated once before stabilizing, and still others intercalated back and forth frequently (Figure 1, Supplementary Data). This highly variable behavior of the central junction is characteristic of complex active systems, in stark contrast to passive systems like foams which maintain relatively stable configurations unless external forces are applied, with the exception of slow aging processes and thermal fluctuations.

### 2.2 Central junction length anticorrelates with central junction tension

Having observed spontaneous intercalations in our system, we next sought to study their regulation. Since junctional dynamics are clearly at the core of the process of intercalation, we wanted to understand its relationship with the related cell-generated forces. We started by investigating the tensions exerted along the contact interfaces. Relative cell-cell interfacial tensions can be accurately inferred from cell geometry, applying a well-established method of force inference from imaging data [22, 21]. By measuring contact angles at vertices (Figure 2A) and applying force balance equations, we calculated relative tensions for the central junction *T_cj_*and lateral junctions *T_l_* (Figure 2B) (see Material and Methods for details). This analysis (Figure 2A) revealed a strong negative correlation between the relative tension of the central junction *T_cj_*and its length (Figure 2C-D, Movie S7). Specifically, as the central junction length decreased, *T_cj_* showed a marked increase, reflecting a tension build-up during junction shrinkage along the junction relative to the lateral contacts.

**Figure 2:**
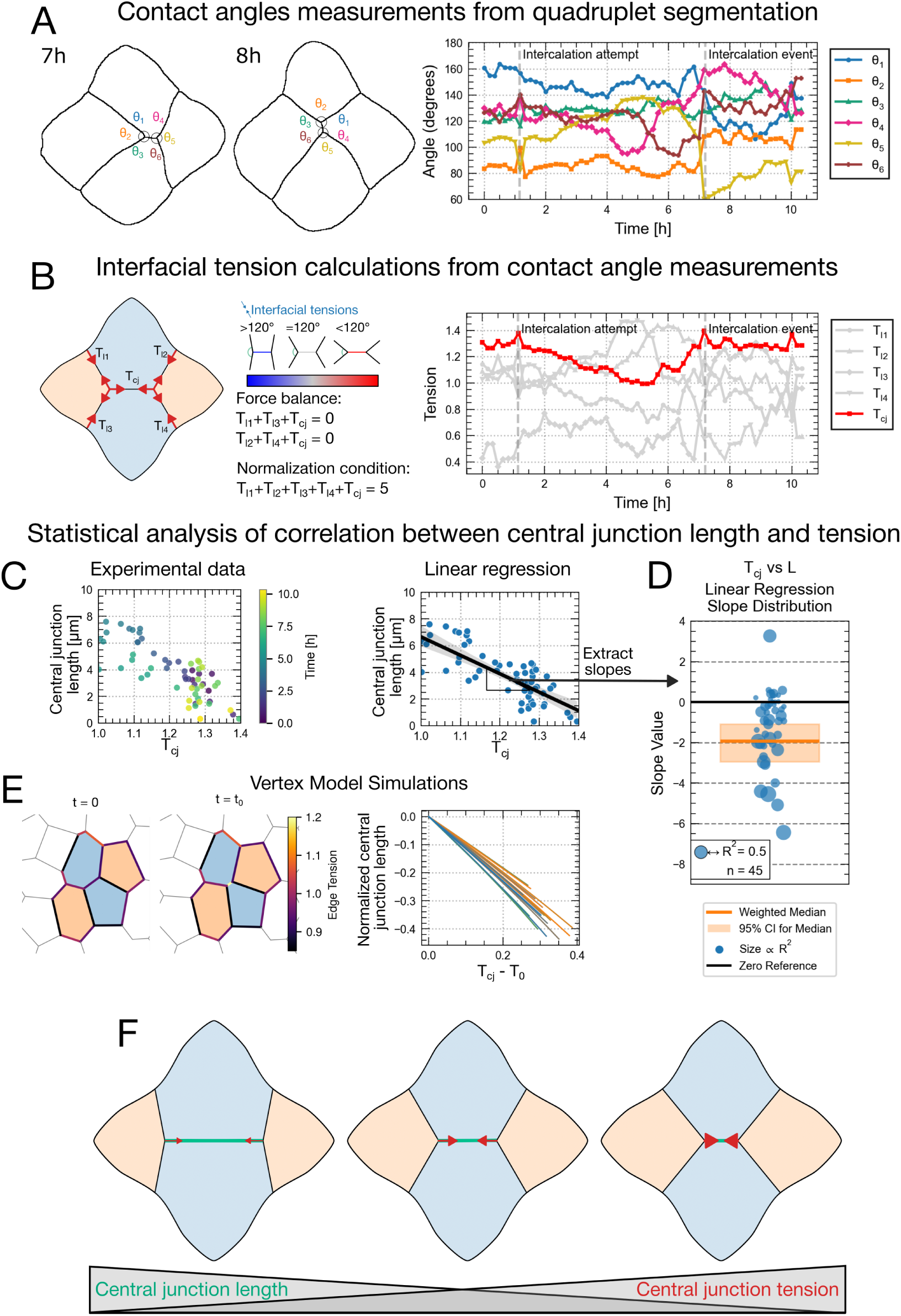
Central junction length anticorrelates with central junction tension. (A) Contact angle measurements in quadruplet cells. Left panel shows segmented cell boundaries at 1h and 3h with labeled angles (*σ*_1_-*σ*_6_). Right panel shows the temporal evolution of all contact angles throughout an intercalation event (marked by vertical dashed line). (B) Calculation of interfacial tensions from contact angle measurements. Left panel illustrates the tension network in the quadruplet, with force balance equations and normalization condition. Center diagram shows the relationship between interfacial tensions and contact angles. Right panel displays the temporal evolution of calculated tensions before and after intercalation. (C) Experimental measurements demonstrating the negative correlation between central junction length and tension (*T_cj_*) with linear regression fit. (D) Statistical analysis of the relationship between central junction tension and length across 45 quadruplets. Plot shows the distribution of linear regression slopes with weighted median (weighted by R^2^) (orange line), 95% confidence interval (shaded area), and individual data points sized according to R^2^ values. (E) Vertex model simulation of junction dynamics. Left panel shows the initial state with color-coded edge tensions, and middle panel shows the configuration after junction shrinking. Right plot shows simulation results showing the negative correlation between normalized central junction length and tension.(F) Schematic summary illustrating the inverse relationship between central junction length and tension, where shorter junctions experience higher relative tension. The relationship between central junction length and relative tension over time is illustrated in Movie S7.

Analysis of the slope of these correlations across multiple quadruplets confirmed robust relationships between *T_cj_* and central junction length. Figure 2C shows this correlation for an example and Figure 2D shows the aggregated slope and correlation strength data. Vertex model simulation provided additional insight into the tension dynamics of the quadruplet system [27, 28, 29]. These simulations (Figure 2E) reproduced the observed inverse relationship between central junction tension and length, further validating the experimental observations.

Figure 2E shows a schematic summary of this conclusion. These results both corroborate existing data from drosophila [11, 30, 16, 31] and validate the relevance of our quadruplet assay.

### 2.3 Central junction length correlates with migratory activity

Since migratory activity has been proposed to be a key driver of cell intercalation [17, 18, 14], we next sought to measure its contribution in our quadruplet system. We combined actin live imaging with traction force microscopy (TFM) to map the spatial distribution of forces the cells exerted on the underlying substrate, while simultaneously tracking the length of the central junction (Figure 3).

**Figure 3:**
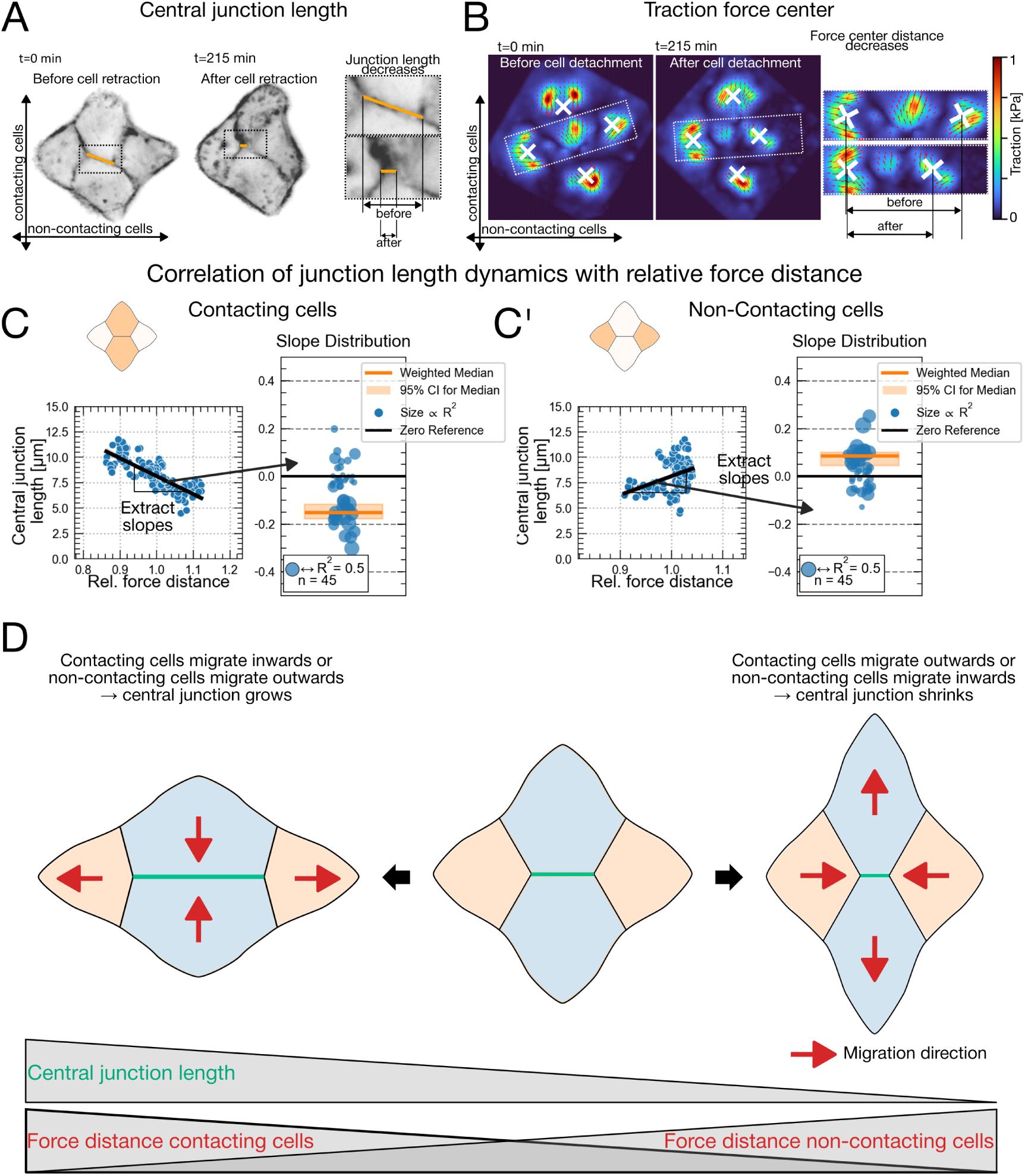
Central junction length correlates with migratory activity. (A) Central junction length changes before and after cell retraction. Left images show actin-labeled quadruplet at t=0 and t=215 min with central junction highlighted in orange. Right panel shows enlarged view comparing junction length before and after detachment. (B) Traction force maps of the same quadruplet before and after cell retraction. Left panels show spatial distribution of traction forces (color scale in kPa). Right panel illustrates the decreased distance between force centers following detachment. (C) Analysis of relationship between central junction length and relative force distance between contacting cells. Left panel shows correlation plot with linear regression line for an example quadruplet. Right panel displays distribution of regression slopes across 45 quadruplets, with weighted median (orange line), 95% confidence interval (shaded area), and individual data points sized according to R^2^values (median R^2^=0.5). (C’) Similar analysis showing relationship between central junction length and relative force distance between non-contacting cells, revealing a negative correlation. (D) Schematic summary of how migratory activity influence central junction dynamics. When contacting cells migrate inward, the central junction grows; when they migrate outward, it shrinks. Conversely, when non-contacting cells migrate outward, the central junction grows; when they migrate inward, it shrinks. The correlation between traction force centers and junction dynamics is illustrated in Movie S8.

Since direct migratory forces are transient and difficult to measure, we used the centers of these traction forces as a measurable proxy for migratory activity (see Materials and Methods for details). This approach revealed a striking relationship between the distance separating the force centers and junction dynamics. We observed, for example, that cell retraction events corresponded with significant changes in both central junction length and the traction force distribution (Figure 3A-B)

Statistical analysis revealed that central junction length positively correlates with the relative distance between force centers of non-contacting cells (Figure 3C, Movie S8), while negatively correlating with the distance between contacting cells’ force centers (Figure 3D, Movie S8). Specifically, when contacting cells migrate toward each other, or when non-contacting cells migrate away from each other, the central junction grows; conversely, outward migration of contacting cells or inward migration of non-contacting cells leads to junction shrinkage (Figure 3E). These findings demonstrate that migratory forces significantly influence junction dynamics even in the absence of directed external cues.

These results suggest that differential cell migration serves as a key driver of spontaneous intercalation, complementing the junctional tension mechanisms described in Figure 2. Our minimal quadruplet system thus reveals how the interplay between cortical tension and cell migration forces collectively regulates cell intercalation in epithelial tissues.

### 2.4 Intercalation mechanics in primary mesoderm quadruplets are similar to those in epithelial cells

To investigate the applicability of our quadruplet approach to developmentally relevant contexts, we adapted the system for Xenopus laevis mesoderm cells (Figure 4). Mesoderm collective internalisation during gastrulation is a core morphogenetic process of early embryogenesis, which crucially relies on extensive intercalations [17, 18]. Rather than isolating native mesoderm, we optimized cell yield and quality through ectopic mesoderm induction by expression of *β*-catenin and constitutively active Activin receptor (caActR) [32, 33].

**Figure 4:**
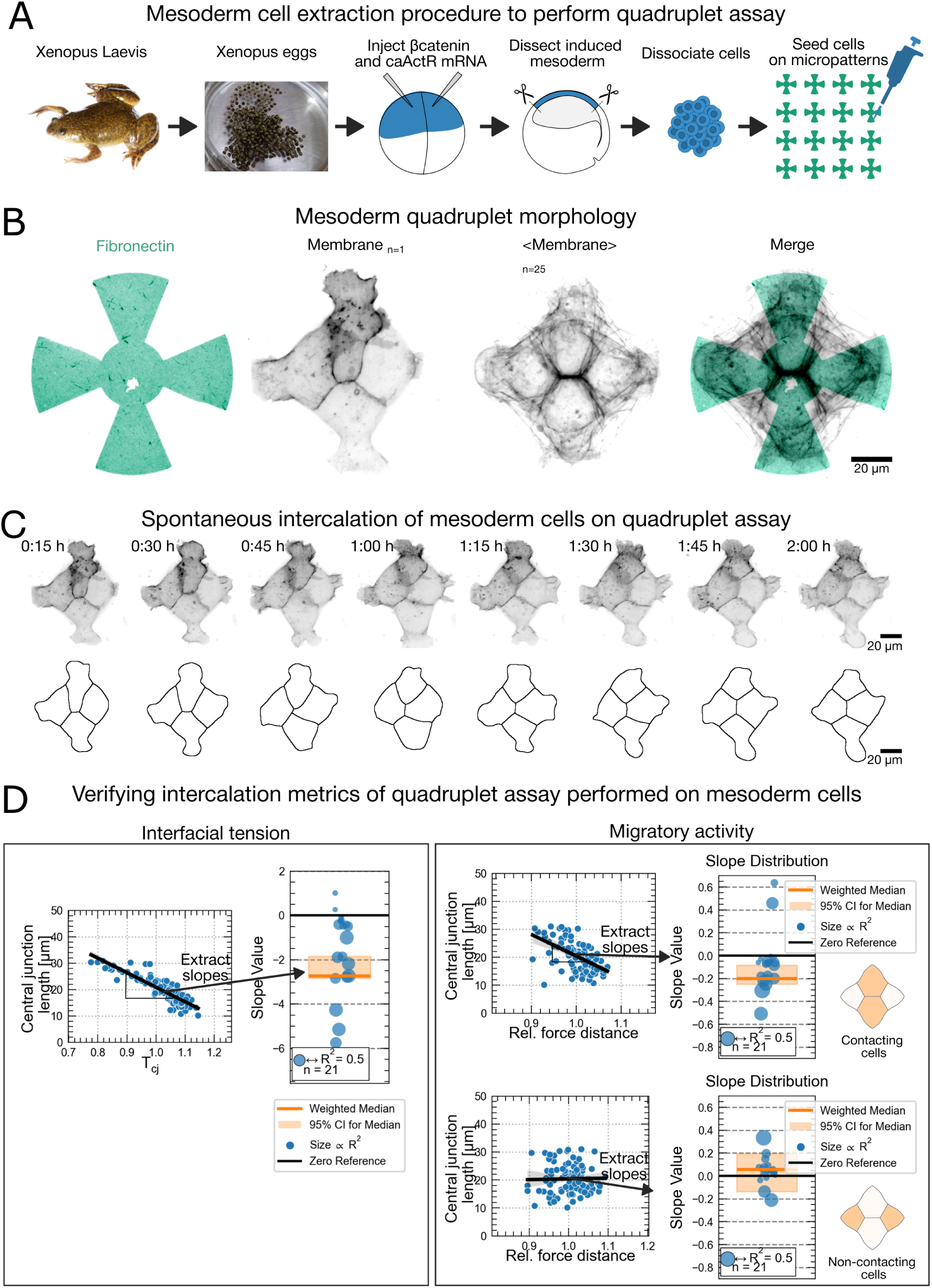
Intercalation mechanics in primary mesoderm quadruplets are similar to those in epithelial cells. (A) Schematic of mesoderm cell extraction procedure. Adult Xenopus laevis frogs provide the eggs that are fertilized and then microinjected with *β*-catenin and caActR mRNA to ectopically induce mesoderm formation in the animal cap. The induced mesoderm is dissected, dissociated into single cells, and seeded onto micropatterned substrates. (B) Mesoderm quadruplet morphology. From left to right: fibronectin windmill-pattern (green); membrane visualization of a representative mesoderm quadruplet; averaged membrane signal from 25 quadruplets showing consistent cellular arrangement; merged image of fibronectin pattern with averaged membrane signal. (C) Time-lapse imaging of spontaneous intercalation in mesoderm quadruplets. Top row shows membrane-labeled cells over 2 hours with corresponding cell boundary segmentations below. (D) Verification of intercalation mechanics in mesoderm quadruplets. Left panel: Analysis of interfacial tension showing negative correlation between central junction tension (*T_cj_*) and length with slope distribution across 18 quadruplets. Right panels: Migratory activity analysis showing correlations between central junction length and relative force distances for both contacting and non-contacting cells, with corresponding slope distributions. Some examples of Xenopus mesoderm quadruplets are shown in Movie S9.

**Figure 4, Supplementary Data.**
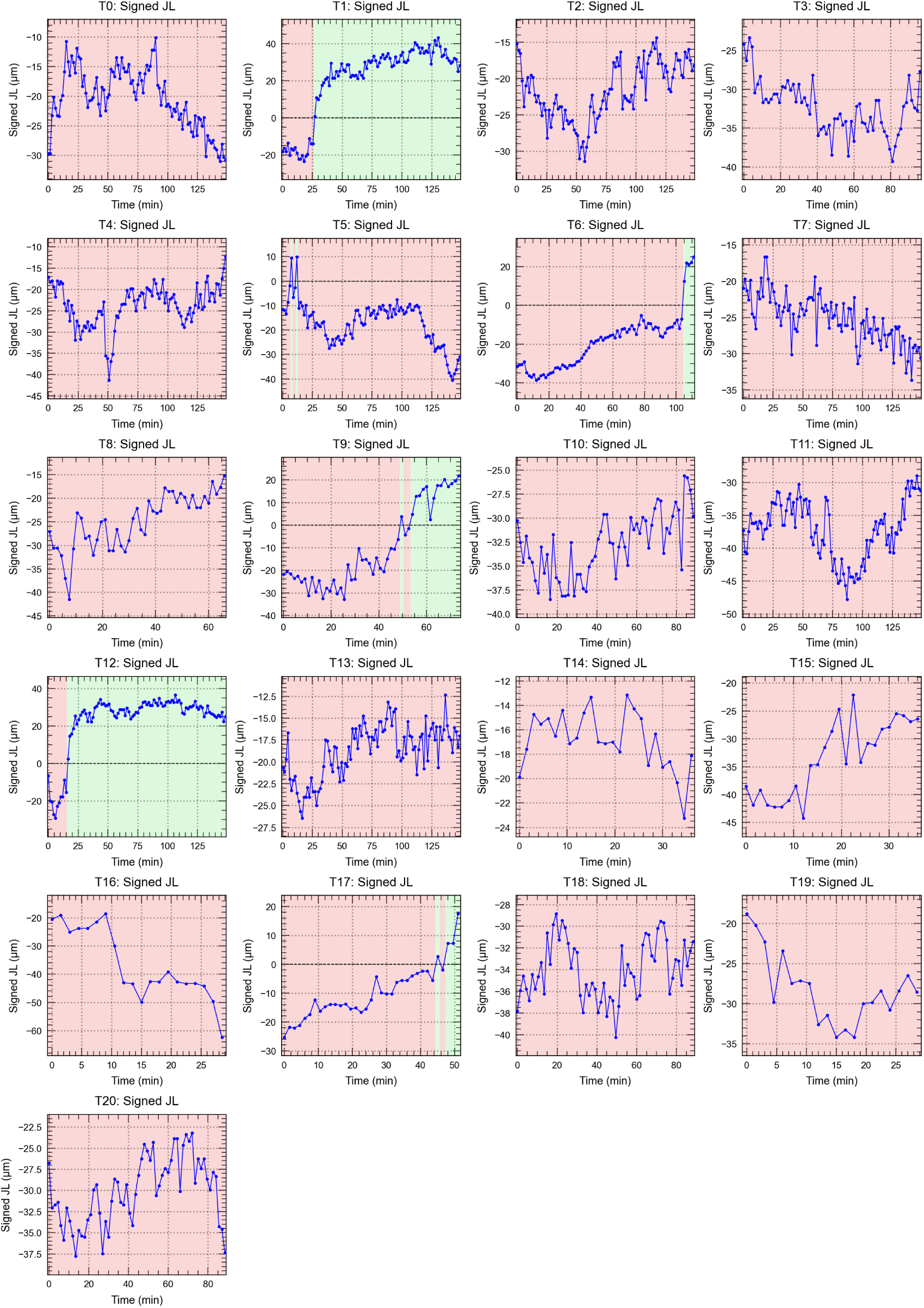
Central junction dynamics and intercalation events in XNPS quadruplets. Signed central junction length over time for individual quadruplets. Negative lengths denote the junction state before an intercalation event, and positive lengths the state after.

Preliminary experiments showed that the mesenchymal-like mesoderm cells required more adhesion area than provided by the cross shape in order to spread fully. To this end, we designed windmill-shaped micropatterns that provided additional spreading area while maintaining the crucial quadruplet geometry (Figure 4B). Live imaging revealed that mesoderm cells reliably formed stable quadruplets on these patterns and underwent spontaneous intercalation events comparable to those observed in MDCK cells (Figure 4C, Movie S9). Intercalations were observed in 6 out of 21 (29%) quadruplets (Figure 4 Supplementary Data). Notably, mesoderm cells exhibited substantially faster dynamics than MDCK cells, with all cellular processes appearing to operate approximately an order of magnitude faster, consistent with the rapid tissue reorganization required during gastrulation.

Importantly, quantitative analysis of the mechanical principles governing intercalation in mesoderm quadruplets revealed striking similarities to our findings with epithelial cells. The inverse relationship between central junction tension and length was preserved (Figure 4D, left panels), showing a robust negative correlation. Similarly, the relationships between migratory forces and junction dynamics were maintained, with central junction length showing consistent correlations with relative force distances between both contacting, but no significant correlations for non-contacting cells (Figure 4D, right panels).

These findings demonstrate that fundamental mechanisms driving intercalation, the interplay between junctional tension and differential cell migration, are conserved across these distinct cell types despite their different developmental origins and morphological and mechanical characteristics. The successful adaptation of our quadruplet assay to mesoderm cells establishes its versatility as a platform for studying the basic principles of cell intercalation across diverse biological contexts relevant to development and disease.

### 2.5 Analysis of effective energy barriers reveals shared physical principles of intercalation across cell types

The preceding analyses of junctional tension and migratory forces were built upon a standardized data pipeline that systematically quantified junction length dynamics over time (summarized in Figure 5A-B). We next leveraged this extensive dataset to probe the underlying physics of intercalation from a thermodynamic standpoint. By aggregating thousands of junction length measurements for both MDCK and XNPS cells, we generated probability distributions for observing a given junction length, P(L). Assuming a Boltzmann-like relationship (*E*(*L*) ln(*P* (*L*))), these distributions were converted into effective energy profiles (Figure 5C). The resulting experimental profiles for both cell types closely resembled theoretical predictions from vertex model simulations, featuring a distinct energy barrier at zero junction length that corresponds to the T1 transition point (Figure 5D).

**Figure 5:**
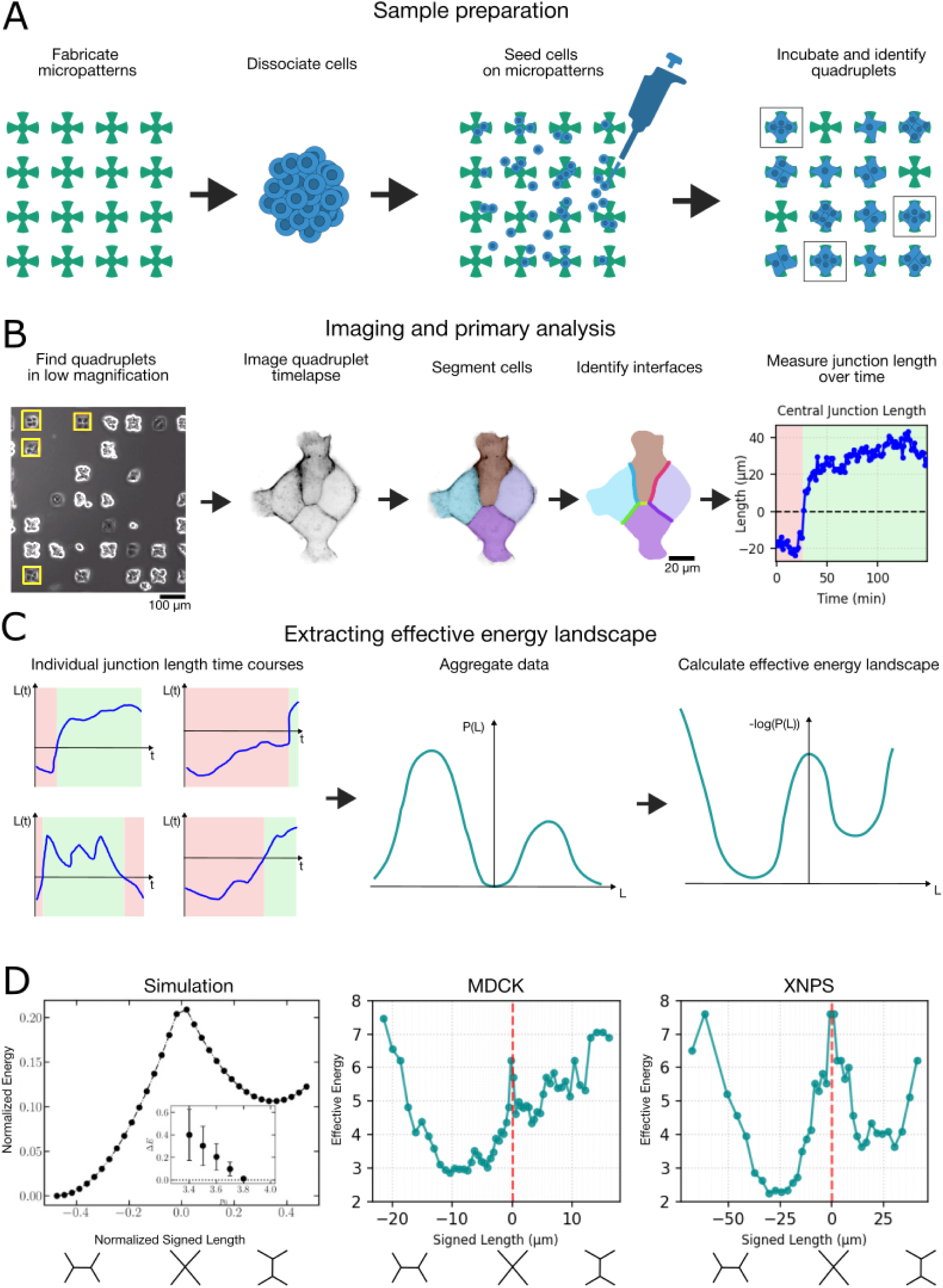
Analysis of effective energy barriers reveals shared physical principles of intercalation across cell types. (A) Schematic representation of the sample preparation procedure, from micropattern fabrication and cell dissociation, through seeding on micropatterns. (B) Imaging and primary analysis workflow showing quadruplet identification, time-lapse imaging, cell segmentation, interface identification, and junction length measurement over time. The pipeline allows extraction of signed junction lengths, where negative values indicate the pre-intercalation state and positive values the post-intercalation configuration. (C) Method for extracting effective energy landscapes from experimental junction length probability distributions. Individual junction length time courses are aggregated to generate probability distributions P(L), which are then converted to effective energy profiles using the relationship −log(P(L)). (D) Effective energy profiles derived from experimental and theoretical data. Left: Theoretical energy profile from vertex model simulations showing characteristic double-well potential with energy barrier at junction length L=0. Middle and right: Experimentally derived effective energy profiles for MDCK and XNPS cells, respectively, obtained by converting normalized junction length probability distributions to effective energies. Both experimental systems show cusp-like barrier profiles similar to theoretical predictions, with dashed vertical lines indicating zero junction length corresponding to the intercalation transition point. Cell configuration diagrams below each panel illustrate the pre-intercalation, transition, and post-intercalation states.

**Figure 5, Supplementary Data 1.**
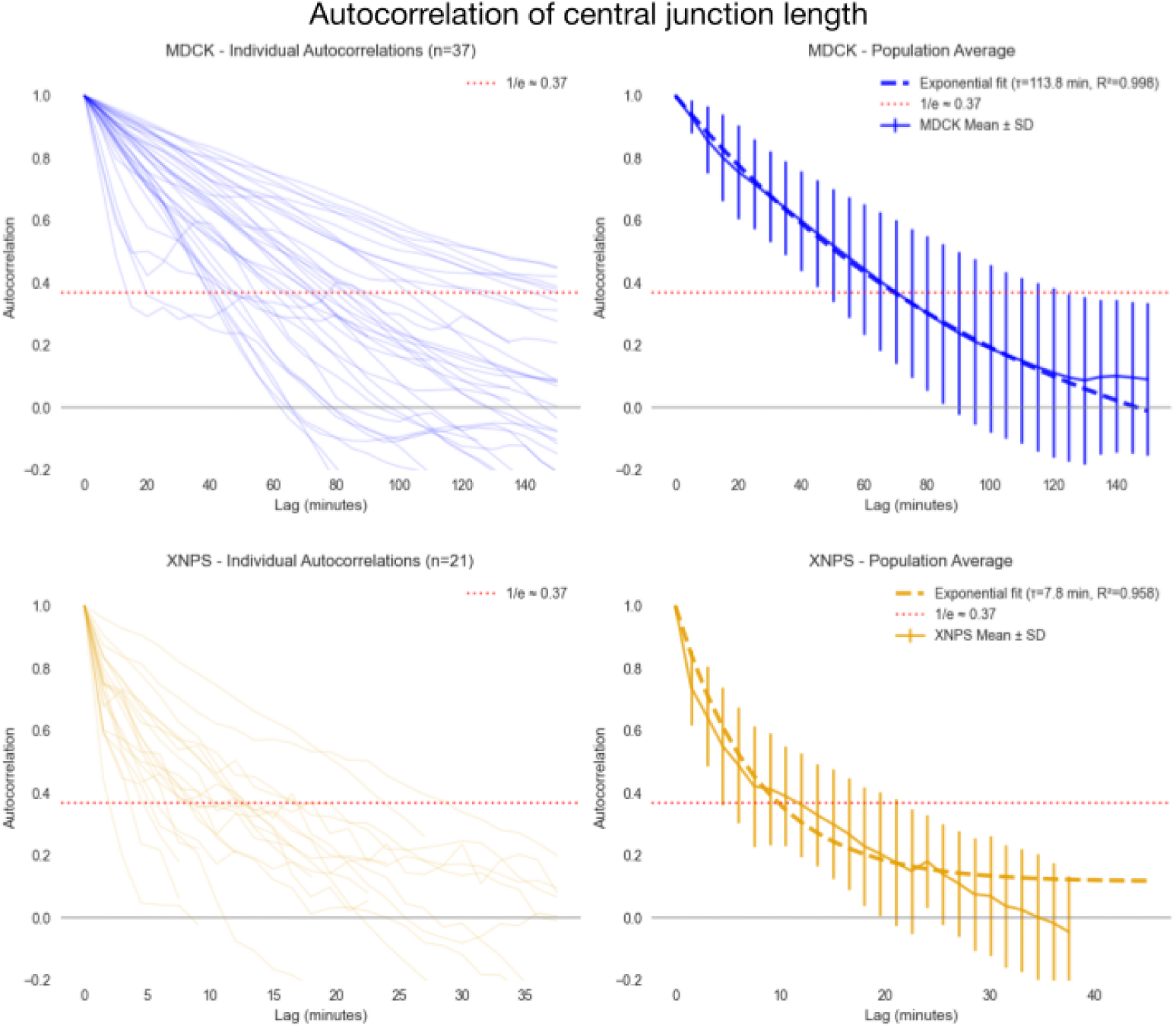
Autocorrelation analysis reveals persistent junction length dynamics in both cell types. Left panels show individual autocorrelation curves for each quadruplet (n=37 for MDCK, n=21 for XNPS), with the dotted horizontal line indicating the 1/e threshold (0.37). Right panels display population averages (solid lines) with standard deviations (error bars) and exponential fits (dashed lines). MDCK cells exhibit a characteristic persistence time of *τ* =113.8 minutes, while XNPS cells show a persistence time of *τ* =7.8 minutes.

**Figure 5, Supplementary Data 2.**
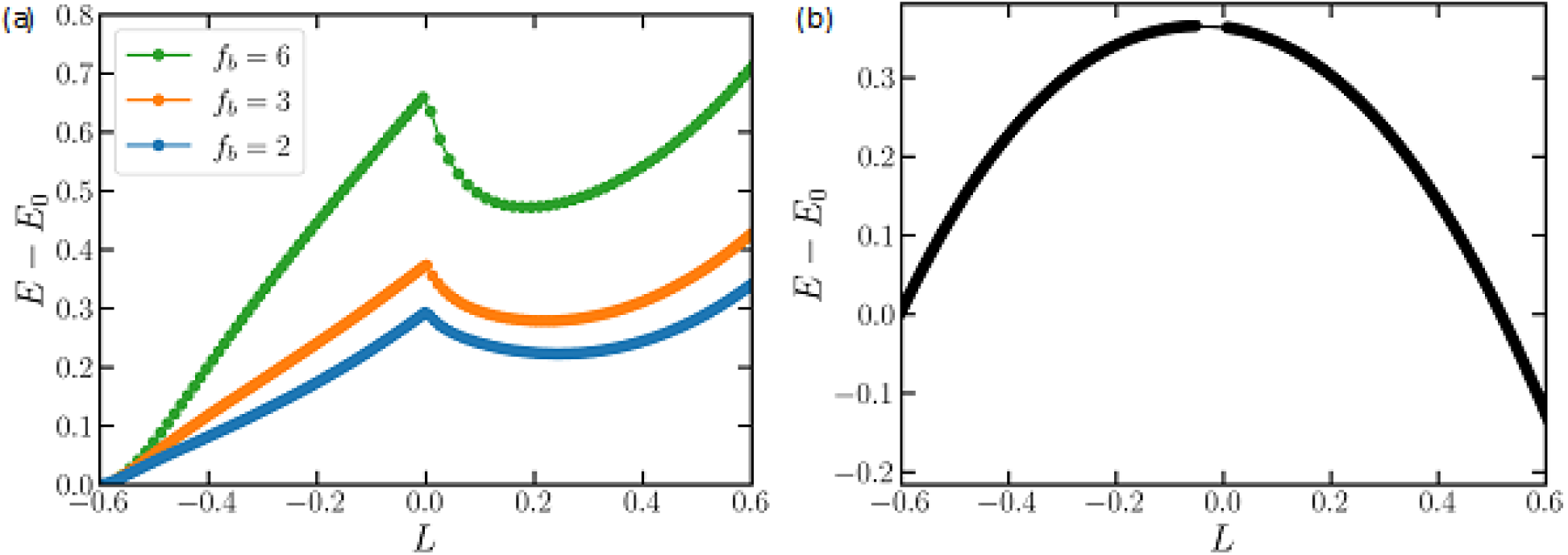
Mechanical barrier of actively driven T1 transitions. (a) Energy difference as a function of edge length for various magnitudes of body forces applied to the non-contacting cells, illustrating how active forces modifies the mechanical barrier. (b) Energy difference for a vertex model incorporating active edge tension, based on the elastic energy in Equation (2).

Despite their different developmental origins and mechanical properties, both epithelial MDCK and mesenchymal XNPS cells exhibited comparable barrier profiles with characteristic cusp-like shapes, suggesting conserved physical constraints governing the intercalation process. However, autocorrelation analysis of junction length dynamics revealed significant temporal persistence in both cell types, with characteristic timescales of approximately 114 minutes for MDCK cells and 7.8 minutes for XNPS cells (Figure 5 Supplementary Data 1). This persistent motion indicates that junction dynamics are actively driven rather than purely stochastic, which challenges the thermal equilibrium assumption underlying the Boltzmann relationship used to derive these energy profiles.

Vertex model simulations incorporating active forces help interpret these findings (Figure 5 Supplementary Data 2). Simulations with active body forces (mimicking migratory activity) preserve the cusp-like energy barrier shape, while simulations with actively ramping edge tensions produce more rounded profiles. The cusp-like profiles observed experimentally are therefore consistent with intercalation being driven primarily by migratory forces rather than active contractile tensions, supporting our traction force analysis.

## 3 Discussion

Our study introduces a novel minimal in vitro assay that bridges the gap between single-cell behaviors and complex tissue dynamics, providing a platform to address critical knowledge gaps in our understanding of cell intercalation. By reducing the intercalation process to its fundamental unit, the four-cell quadruplet, this system enables precise investigation of mechanisms that remain obscured in complex tissue environments. This methodological advancement establishes a controlled experimental framework that can be used to dissect the elementary mechanical principles governing cellular rearrangements during morphogenesis, offering unprecedented opportunities to isolate and quantify the forces driving intercalation events.

### 3.1 Spontaneous intercalation in a minimal system

Perhaps the most striking finding of our work is that such a highly constrained system spontaneously displays intercalation events with remarkable frequency. This was far from intuitive, as one might expect that the geometric constraints imposed by micropatterning would suppress the dynamic cellular behaviors observed in vivo. Instead, our quadruplet system demonstrates that interfacial tension dynamics and differential cell migration are sufficiently robust to operate even under artificial boundary conditions. This suggests that intercalation is an intrinsic property of cell collectives that emerges from local mechanical interactions rather than requiring complex tissue-level coordination.

The spontaneous nature of intercalations in our system is particularly significant given that most previous studies have focused on directionally biased intercalations driven by external morphogenetic cues [11, 34, 19]. However, unbiased intercalations are prevalent in numerous developmental contexts, including mesoderm invagination and intestinal morphogenesis, yet have received comparatively little experimental attention. Our assay provides the first systematic platform for investigating these spontaneous rearrangements under controlled conditions, filling a crucial gap in the field.

### 3.2 Isolating intercalation from tissue-level complexity

Beyond demonstrating that intercalation can occur spontaneously, a key advantage of our minimal system is its ability to uncouple the elementary intercalation event from the complex mechanical environment of a larger tissue. A major limitation of studying intercalation in whole tissues is the presence of cascade effects, where individual T1 transitions influence neighboring cells and propagate mechanical perturbations across the tissue [23, 24]. While it was generally anticipated that intercalation could occur through local mechanical interactions [35], this had not been directly demonstrated in isolation from tissue-level influences. Our findings provide clear evidence that isolated cell quadruplets can undergo intercalation autonomously, driven solely by local mechanical dynamics. This confirms that the elementary unit of tissue rearrangement is mechanically self-sufficient and establishes a foundation for understanding how local intercalations contribute to global tissue deformations.

This isolation of intercalation events enables precise quantitative analysis that would be impossible in complex tissue environments. By removing the confounding effects of long-range mechanical coupling and morphogenetic gradients [36, 37], our system allows direct measurement of the relationship between local migratory forces, interfacial tensions and junction dynamics.

### 3.3 Bridging cell-matrix and cell-cell interactions

Our integration of micropatterning with traction force microscopy provides the first direct quantification of both cell-matrix and cell-cell forces during intercalation. While the unified model of intercalation has established that cell migration and interfacial tension work together [10], previous studies have characterized migratory activity through imaging of molecular markers such as myosin and actin within embryonic tissues, where direct force measurement remains limited and technically challenging. Our approach enables simultaneous measurement of traction forces exerted on the substrate and quantification of interfacial tensions, providing direct mechanical validation that migratory forces play a crucial role in junction dynamics alongside the well-established importance of interfacial contractility [11, 30, 16, 31].

### 3.4 Energy landscapes and active matter physics

The field of active matter physics has developed sophisticated theoretical frameworks for understanding non-equilibrium biological systems [38], and our experimental platform provides an ideal system for testing these theories. Our novel approach of deriving effective energy profiles from junction length probability distributions using 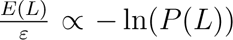 represents a significant methodological advance, yet reveals fundamental challenges in applying equilibrium concepts to active cellular processes.

This energy landscape methodology relies on the assumption of thermal equilibrium, where the probability of observing a particular configuration is governed by its energy through the Boltzmann distribution. However, our autocorrelation analysis clearly demonstrates that junction dynamics exhibit substantial temporal persistence, with characteristic timescales ranging from approximately 8 minutes for XNPS cells to 114 minutes for MDCK cells. This persistent motion indicates that cells actively drive intercalation through directed forces rather than only through fluctuations around equilibrium states, fundamentally challenging the equilibrium assumption underlying our energy landscape interpretation.

This apparent contradiction between the active nature of cellular systems and equilibrium-based theoretical frameworks creates a productive tension that highlights the need for active matter approaches. Our vertex model simulations provide crucial insights into this relationship, showing that simulations incorporating active body forces (mimicking cell migration) preserve the cusp-like barrier shape observed experimentally, while actively ramping edge tensions produce more rounded profiles. The cusp-like profiles observed experimentally are therefore consistent with intercalation being driven primarily by migratory forces rather than active contractile tensions. This suggests that the shape of experimental energy profiles may serve as a diagnostic tool for identifying the dominant active mechanisms driving intercalation.

The framework we have established provides a powerful platform for resolving these theoretical questions through systematic experimental investigation. Future studies can leverage our quadruplet system to test predictions from active matter theory by systematically varying the strength and nature of active forces through optogenetic perturbations, pharmacological treatments, or mechanical manipulations. The high temporal resolution and quantitative nature of our measurements make it possible to distinguish between different theoretical models and validate emerging frameworks for understanding active biological systems.

### 3.5 Translational potential and accessibility

Finally, our demonstration that this assay can be extended across different cell types establishes its broad applicability for developmental biology research. Importantly, while our comprehensive analysis benefited from traction force microscopy, the core readout, the central junction length dynamics, can be measured using standard fluorescence microscopy available in most laboratories. This accessibility makes the quadruplet assay a practical tool for screening the effects of genetic perturbations, pharmacological treatments, or environmental conditions on intercalation dynamics.

We envision this system becoming a standard assay for the developmental biology community. The simplicity of the readout, combined with the quantitative nature of the measurements, makes it ideal for high-throughput screening approaches. We are committed to sharing our micropatterning protocols, analysis software, and expertise with laboratories interested in adopting this approach.

### 3.6 Future directions and broader implications

Looking forward, our quadruplet system opens numerous research directions. The ability to precisely control initial cell configurations and systematically vary pattern geometries provides unprecedented opportunities to dissect how local tissue architecture influences intercalation dynamics. Furthermore, the system’s compatibility with advanced imaging techniques and optogenetic perturbations positions it as a powerful platform for investigating the molecular mechanisms underlying the mechanical behaviors we have characterized.

In conclusion, our quadruplet assay represents both a significant methodological contribution and experimental validation of existing models for intercalation mechanics. By demonstrating that fundamental tissue behaviors can emerge from minimal cellular assemblies, we provide direct mechanical evidence supporting the relationship between local cellular interactions and global tissue dynamics. The coupling of our assay with vertex modeling further suggests a dominant role for migratory forces in driving the observed intercalation events. This work establishes a foundation for future investigations into the physical principles governing morphogenesis and offers practical tools for the broader developmental biology community.

## 4 Materials and Methods

### 4.1 Constructs

Plasmids used in this study were a constitutively active Alk-4 (activin receptor type 1-B) in pSP64T, Myc-tagged full length Xenopus *β*-catenin and membrane GFP, both in pCS2+ vector. Capped mRNAs were synthesized according to manufacturer instructions (mMessage mMachine kit, Ambion).

### 4.2 Cell Culture

MDCK cells were cultured in DMEM supplemented with 10% FBS and 1% penicillin/streptomycin at 37°C and 5% CO_2_. For experiments using MDCK quadruplets, cells were seeded at a density of approximately 30,000 cells per coverslip and allowed to attach and divide for 16-28 hours.

### 4.3 Xenopus Embryo Manipulation and Mesoderm Induction

For Xenopus mesodermal cells, embryos were obtained by in vitro fertilization and staged according to Nieuwkoop and Faber [39]. To optimize cell quality and yield, we used an established protocol for ectopic mesoderm induction rather than extracting native mesoderm [32, 33]. This approach allowed us to generate large numbers of mesodermal cells suitable for micropatterning and imaging experiments. Induced mesoderm offered several advantages over native mesoderm extraction: it produced substantially higher cell numbers required for micropatterning experiments, and the cells have less yolk content resulting in improved optical transparency for high-resolution imaging. To induce mesodermal cell fate, a mixture of mRNA encoding *β*-catenin (500-100 pg) and constitutively active Activin receptor (caActR, 200-1000 pg) were injected into the animal hemisphere of the two blastomeres of 2-cell stage embryos [32, 33]. At stage 10 (early gastrula), the animal cap tissue was dissected in 1× MBSH (88 mM NaCl, 1mM KCl, 2.4 mM NaHCO_3_, 0.82 mM MgSO_4_, 0.33 mM Ca(NO_3_)_2_, 0.33 mM CaCl_2_, 10 mM Hepes) and dissociated into single cells by incubation in calcium-free alkaline buffer (88 mM NaCl, 1 mM KCl and 10 mM NaHCO_3_, pH = 9.5) [40]. Cells were seeded on the micropatterns in 1× MBSH.

### 4.4 Micropatterning

For MDCK cells, micropatterned substrates were prepared using deep UV photolithography as described in [41, 26]. Briefly, glass coverslips were cleaned with plasma and coated with PLL-g-PEG (0.1 mg/mL)(SuSoS, catalog name: PLL(20)-g[3.5]-PEG(5)). A photomask containing the desired patterns was placed on the coverslip and exposed to deep UV light for 5 minutes. The exposed areas were then coated with fibronectin (20 *µ*g/mL) (Sigma F1141) and fluorescently labeled fibrinogen (20 *µ*g/ml)(Sigma F13192).

For XNPS cells, microcontact printing was used. PDMS stamps with the windmill pattern were cast from silicon masters, coated with fibronectin solution (50 *µ*g/mL) (Sigma F1141) and fibrinogen solution (50 *µ*g/mL)(Sigma F13192), and brought into contact with glass coverslips for 5 minutes. Silicon masters were prepared by photolithography of SU-8 on silicon wafers [42].

### 4.5 Traction Force Microscopy

Polyacrylamide gels (Young’s modulus = 20 kPa for MDCK, 6 kPa for XNPS cells, thickness ≈ 50 *µ*m) were prepared by mixing acrylamide/bis-acrylamide solutions with fluorescent beads (Thermo Fisher F8807). The gels were functionalized with the micropatterned protein using either photolithography or microcontact printing methods described above.

Cell-generated forces were measured using batchTFM, an open-source Python tool for traction force microscopy [26]. Bead displacements were tracked using optical flow (TV-L1 algorithm), and traction forces were calculated using Fourier Transform Traction Cytometry with a regularization parameter determined empirically. The analysis pipeline includes drift correction, contrast enhancement, and force reconstruction validated using synthetic data sets.

### 4.6 Image Acquisition

Time-lapse imaging of MDCK cells was performed on a Nikon Ti2-E microscope equipped with a Perfect Focus System, temperature control, and CO_2_ regulation. Images were acquired using a 60× objective (NA 1.4). For live imaging of filamentous actin, cells were stably transfected with far-red LifeAct. All imaging was performed under controlled environmental conditions with appropriate temperature and CO_2_ regulation to maintain cell viability throughout the experimental duration.

Experiments on Xenopus cells were performed at room temperature on an Olympus IX83 inverted microscope coupled to a Yokogawa W1 spinning disk unit, including a SoRa disk and a Z-drift correction system using a 60× objective (NA 1.3). For membrane visualization, cells were labeled with membrane GFP.

### 4.7 Segmentation and Cell Interface Analysis

Cell segmentation and cell interface analysis was performed using our custom-developed napari plugin, napariCellFlow, which incorporates Cellpose 2.0, a deep learning-based segmentation algorithm for robust cell identification [43, 44]. The napariCellFlow plugin provides a comprehensive analysis pipeline including cell segmentation, cell tracking, edge detection, and intercalation event analysis.

For cell segmentation, we used the Cyto2 as base model from Cellpose. After segmenting a few images, we used these to train a custom model based on Cyto2. Cell diameter was set based on visual inspection using scale disk overlays, with flow threshold and cell probability parameters adjusted to achieve optimal cell separation and detection confidence.

Cell tracking across time frames was performed using napariCellFlow’s custom tracking algorithms.

Edge and intercalation detection were implemented using custom algorithms within napariCellFlow. The plugin automatically detects cell boundaries, measures junction lengths, and identifies T1 transition events based on topological changes in cell connectivity. Junction length measurements were extracted by analyzing the contact interfaces between neighboring cells.

The resulting segmentation masks and tracking data were used to quantify interface geometry for force analysis, and junction length dynamics over time. Force measurements were analyzed using custom Python scripts, with statistical analysis performed using scipy and graphs generated using matplotlib.

The napariCellFlow code is available at https://github.com/ArturRuppel/napariCellFlow under the GPL-3.0 license, providing full reproducibility and accessibility for the developmental biology community.

### 4.8 Displacement of Cell Center of Forces

We observed that the central junction length between contacting cells correlates with the migration of non-contacting cells. To model this behavior, we employ an overdamped system where the migration of cells is driven by a pulse of migratory force *F* acting on each cell. The position *x*(*t*) of the cell evolves under the balance of this force and a substrate friction. Given that the motion is overdamped, the forces balance the friction at each time step, leading to the following equation for the velocity:

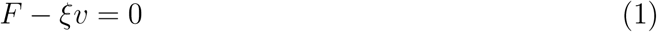

This can be integrated over time to give the displacement of the cell from its initial position *x_i_*. The final position *x_f_* after a time *t* is:

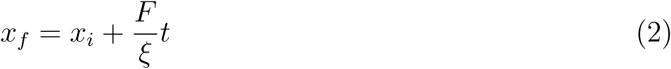

Here, 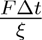 is the migration-driven displacement, which scales with the force and the friction coefficient of the substrate. Experimentally, we cannot directly measure this pulse of migratory force acting on these cells. Thus, we measure the distance between the centers of traction forces on the non-contacting cells and find that this distance increases as these cells migrate away. This is associated with an increase in the central junction length between contacting cells. In contrast, the decrease in the vertical (y-axis) distance between the traction centers of the contacting cells is correlated with a reduction in the junction length, suggesting that as the contacting cells migrate away, there will be space for the non-contacting cells to come closer, hence shortening the junction. Thus, the overall migration process of non-contacting cells, modeled by this simple force-displacement relation, explains the observed behavior of junction length changes. The traction forces provide the necessary *F*, while the time evolution and friction *ξ* describe the dynamics of these changes in junction length.

### 4.9 Vertex Model

The vertex model provides a coarse-grained framework to study the mechanics of epithelial tissues by representing them as networks of polygonal cells. These cells are generated by performing a Voronoi tessellation of randomly placed seed points within a two-dimensional box with periodic boundary conditions. The tessellation ensures that the initial configuration consists of polygons representing cells, which share edges with neighboring polygons. The vertices of these polygons serve as the physical degrees of freedom of the model.

The mechanics of the tissue are governed by an energy functional that incorporates key physical and biological properties of cells. The energy for a system of *N* cells is expressed as:

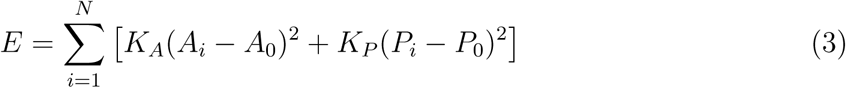

where *A_i_* and *P_i_* are the area and perimeter of cell *i*, respectively. The parameters *A*_0_ and *P*_0_ represent the preferred area and perimeter of the cell. The area term penalizes deviations from the preferred area, reflecting cell incompressibility and resistance to height fluctuations, while the perimeter term captures active contractility of the actomyosin cortex and the balance of cortical tension with cell-cell adhesion. The coefficients *K_A_* and *K_P_*are stiffness parameters that quantify the relative importance of these contributions. The energy of the system is minimized with respect to the cell vertex positions to determine the mechanical equilibrium configuration of the tissue.

A crucial dimensionless parameter used to characterize mechanics of vertex models is the preferred cell shape factor *p*_0_ = *P*_0_*/√A*_0_. Tissues with a preferred cell shape factor below a critical value *p*^∗^ ≈ 3.8 are typically rigid-like, while those with *p*_0_ *>* 3.8 exhibit a fluid-like behavior.

To directly simulate the effect of edge tensions on edge length in Fig. 2, we use a modified vertex model energy given by:

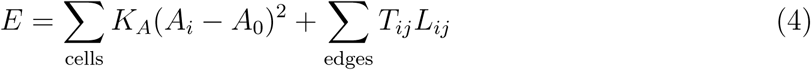

where, instead of modeling perimeter elasticity of cells, we introduce a direct tensional energy term for each edge between neighboring cells. Here, *T_ij_* represents the tension, and *L_ij_* is the length of the edge *ij*. This modification allows us to explicitly capture the effect of active edge tensions in our simulations.

### 4.10 Energy Barriers for T1 Transitions with Edge Length as the Control Parameter

In the vertex model framework, T1 transitions represent fundamental topological rearrangements where a shared edge between two neighboring cells disappears, and a new edge forms between two previously non-adjacent cells. To compute the energy barrier associated with a T1 transition, we focus on a single edge and quasistatically shrink its length while allowing the rest of the tissue to relax according to the energy functional.

The procedure begins by pinning the two vertices at the ends of the edge while incrementally reducing the edge length. At each step, the positions of all other vertices are relaxed to minimize the total energy of the system, ensuring that only the edge being studied is constrained. This process continues until the edge length becomes smaller than the T1 transition threshold defined in the model. At this point, a manual T1 transition is performed: the topology of the four cells surrounding the edge is updated, replacing the contracting edge with a new edge oriented perpendicularly to the original. After the T1 transition, the new edge is gradually extended by moving the two vertices apart, again allowing the rest of the tissue to relax at each step.

The total energy of the tissue is computed as a function of the edge length throughout the entire process. By defining the initial energy before the edge is manipulated as *E*_0_, the energy difference Δ*E* = *E − E*_0_ is plotted against the edge length, with negative values corresponding to the shrinking phase (pre-T1) and positive values corresponding to the extension phase (post-T1). This energy profile typically exhibits a peak at the transition point (edge length = 0), representing the energy barrier for the T1 transition. The energy barrier is quantified as the difference between the peak energy and the initial energy of the tissue. To obtain statistically robust results, this procedure is repeated for multiple edges in a given simulation and across at least ten different random tissue realizations. The mean value of the computed energy barriers is then reported.

This analysis is performed across a range of preferred cell shape factors. As expected, the energy barriers decrease with increasing *p*_0_, vanishing entirely when *p*_0_ ≳ 3.8, where the tissue transitions from a solid-like to a fluid-like state.

### 4.11 Analysis of Junction Length Probability

Understanding the mechanical properties of tissues often requires linking geometric features to energy landscapes. A notable example is the work by Popovíc et al. [24], which analyzed the bond length distribution in the developing Drosophila wing epithelium to infer the material’s plasticity and yielding behavior. By measuring the distribution of short edges in epithelial networks, they established a connection between local bond lengths and the density of weak spots, demonstrating that the observed scaling was consistent with predictions from the vertex model. Building on this idea, we analyze the probability distribution of junction lengths in MDCK and XNPS epithelial tissues to extract the energy barriers associated with T1 transitions. In Popovíc et al. [24], the data was aggregated across many edges in a sample. This means that the edge length distribution they observed may have resulted from stress-based interactions between T1s across the tissue, which is indeed consistent with the scaling laws they observed.

Building on this idea, we analyze the probability distribution of junction lengths in MDCK and XNPS epithelial tissues to extract the energy barriers associated with T1 transitions. Unlike the statistical approach in Popovíc et al. [24], which aggregated data across many edges to test theoretical scaling laws, our approach focuses on a single four-cell junction over time, and therefore explicitly excludes interacting T1s. This allows us to directly compute the probability *P* (*L*) of observing a specific junction length *L* during the tissue’s evolution and infer the corresponding energy landscape.

To estimate the energy barriers associated with T1 transitions in experimental systems, we analyze the distribution of junction lengths shared by four cells in MDCK and XNPS tissues. Unlike vertex model simulations, where junctions can be directly manipulated, we infer energy barriers by assuming a Boltzmann-like relationship between the energy and the probability *P* (*L*) of observing a junction with length *L*:

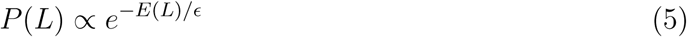

where *ɛ* represents the scale of energy fluctuations. This assumption is based on the principle that in systems exhibiting quasi-equilibrium behavior between successive transitions, the probability of a given configuration is proportional to the exponential of its energy. In densely packed systems, such as epithelial tissues, local rearrangements (T1 transitions) occur when the system overcomes energy barriers separating metastable states. This relationship implies that the energy landscape associated with junction lengths can be expressed as:

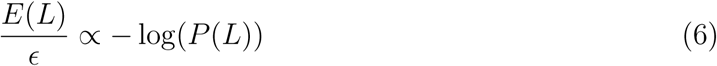

To estimate *P* (*L*), we bin the junction length data for each experimental sample, separately for MDCK and XNPS tissues. The number of data points in each bin is counted, and the probability is computed by normalizing the counts across all bins. The resulting distribution is plotted as − log(*P* (*L*)) versus *L*. Similar to vertex model simulations, we assign negative values to *L* for junction lengths prior to T1 transitions and positive values for lengths after the transition. This analysis yields an energy profile with a peak at *L* = 0, corresponding to the T1 transition. The height of this peak relative to the initial energy provides an estimate of the energy barrier for the transition. Notably, the energy profiles extracted from *P* (*L*) in experiments closely resemble those predicted by our vertex model analysis. This suggests that, despite experimental complexities such as active fluctuations and cell heterogeneity, the fundamental mechanics of epithelial rearrangements align with theoretical expectations.

### 4.12 Actively Driven T1 Transitions

While the above analysis captures *P* (*L*) expected for fluctuations around a passive energy barrier, our observation of changes to traction centers suggest that cells in our quadruplets are subject to directed, active or contractile forces that can drive topological rearrangements. In the main text, we demonstrated that these active traction forces affect the central junction length. To incorporate activity due to pushing and pulling forces, we add body forces to the two non-contacting cells adjacent to the T1 edge (see the simulation movie). We apply the force *f_b_* on these cells, which is technically divided equally between the cell vertices with a magnitude *f_b_/n*, where *n* is the number of cell vertices. Concretely, these active forces are oriented along the edge direction and gradually pull the cells toward the topological transition. We perform an overdamped dynamic simulation with the elastic energy in Eq. (1), applying the active force at each timestep and allowing the vertices to evolve. When the edge length falls below a threshold, we manually carry out the T1 update, then reverse the force direction to extend the new edge.

Even though these body forces push the system out of equilibrium, plotting Δ*E* = *E − E*_0_ versus the edge length still reveals a distinct energy peak at *L* = 0. As shown in Figure S1 (a) for three different force magnitudes *f_b_* in units of *K_A_A*^3*/*2^, the system’s mechanical barrier persists, albeit at different heights due to the stronger deformations imposed by larger active forces. Increasing *f_b_* both raises the maximum of Δ*E* and alters the post-T1 profile, indicating that the tissue experiences more pronounced cell-shape distortions during the transition.

If there is a separation of timescales so that the energy changes generated by directed forces are slow compared to energy changes induced by fluctuations, then fluctuations would explore this effective landscape. This is similar to a material subject to quasi-static shear strain or random active forces [45] or the mechanical loading of a protein that is being stretched [46], where one can study how the energy landscape changes as a function of one reaction coordinate (in this case *L*) while applying a loading or force. Fig 5 Supplementary Data 2 demonstrates the landscape remains cusp-y as a function of *L* in the presence of body forces, which suggests that we should still see a cusp-y *P* (*L*) in experiments where there are active migratory forces, provided that the magnitude of fluctuations are sufficiently large to explore the landscape. Interestingly, our models also predict that if there were no fluctuations (i.e. completely deterministic motion of the system over the energy barrier via *f*_b_), at the transition point there would actually be a peak in *P*(*L*), corresponding to a minimum in – *log*(*P*(*L*)), as the system would decelerate as it traverses uphill in energy and move more slowly through the region at the top of the barrier.

An alternative approach to actively inducing T1 transitions is to modify the elastic energy of the vertex model directly. Instead of applying external body forces, we can drive the transition by gradually increasing the tension on the target edge while minimizing the system’s elastic energy. For example, such active T1s have been identified in germband extension in Drosophila [47]. It is important to emphasize that the actively ramping up tension shrinks the interface even faster than it would shrink under constant large tension, and represents active, directed dynamics of the interface. The total energy of the system is given by Eq. (2). In this method, we slowly increase the tension on the shrinking edge, allowing the system to minimize its energy at each step. Once the edge length falls below a predefined cutoff, the T1 transition is executed, after which the tension is reduced to allow the new edge to extend in the perpendicular direction. The entire process remains quasi-static, ensuring that the system continuously relaxes into local equilibrium. We analyze the energy barrier associated with this tension-driven transition by plotting Δ*E* as a function of the edge length *L*. Similar to the passive and body-force-driven cases, the effective energy landscape exhibits a peak at *L* = 0, confirming the existence of a mechanical barrier. However, the shape of the Δ*E* curve differs significantly from the previous cases. Unlike the sharp cusped profile observed in passive and body-force-driven transitions, the tension-driven energy landscape is smoother and more rounded (Figure 5 Supplementary Data 2 (b)), suggesting that the transition occurs more gradually.

We have checked that the shape of the curves (cusp-y for migratory forces, rounded for actively ramping up tensions) is the same regardless of the energy function (vertex model with a term quadratic in the perimeter *P* vs. foam which has terms only linear in *P*).

## Acknowledgments

We thank Benôıt Charlot and Audrey Sebban for their expert assistance in fabricating silicon masters for microcontact printing. We are grateful to Christine Fagotto-Kaufmann and Guillaume Desgarceaux for sharing Xenopus embryo management responsibilities. We acknowledge the imaging platform MRI (Montpellier Ressources Imagerie) for providing access to microscopy facilities and technical support. We also thank the animal facility ZEFIX (zebrafish and xenopus platform) for maintaining our animal colonies and providing expert care. This work was supported by the ANR (Agence Nationale de la Recherche) grant Inters-cal (ANR-21-CE13-0042), coordinated by François Fagotto. Thomas Boudou acknowledges funding from CNRS grants (PEPS CNRS-INSIS 2021, Lumìere Visible et Vie 2022) and from the ANR CONTRACTILE project, grant ANR-23-CE13-0037 of the French ANR.

